# COCOA.jl – A Julia package for high-performance analysis of concordance and kinetic modules in biochemical networks

**DOI:** 10.64898/2026.05.05.722856

**Authors:** Anton Schaffranke, Anika Küken, Zoran Nikoloski

## Abstract

**Summary:** Recent advances in analysis of biochemical networks have contributed the identification of their modular structure based on the concept of multi reaction dependencies and kinetic coupling of reaction rates (Küken et al., 2022; Langary et al., 2025). Existing implementations of the algorithms to study modular structure do not scale well with the size of the networks, prohibiting their application with genome-scale networks. Here, we introduce COCOA.jl, a multithreaded Julia package for identification of concordant and kinetic modules, with applications in the study of concentration robustness.

**Availability and implementation:** COCOA.jl is implemented in Julia 1.12.2 and is freely available under the MIT license at https://github.com/antoniofranky/COCOA.jl. It runs on Linux, macOS, and Windows; installation is supported via the Julia package manager. COCOA.jl can be called from Python via JuliaCall.

## 1 Introduction

Constraint-based modelling represents a key tool for genome-scale analysis of cellular metabolism, drug-target discovery, and design of metabolic engineering strategies (Bintener et al., 2023; Gu et al., 2019; Thiele et al., 2020). Classical flux coupling analysis characterises pairwise dependencies between individual reactions (Burgard et al., 2004; Larhlimi et al., 2012). Recent theoretical advances have enabled the identification of multireaction dependencies and linked them to concentration robustness in biochemical networks endowed with mass-action kinetics.

These methods operate on the complex–reaction decomposition of the stoichiometric matrix, **N = YA**, where each complex (one side of a reaction) carries an activity equal to the net flux through it. Complexes with zero activity across all feasible steady states are called balanced (Küken et al., 2021; Langary et al., 2023); two non-balanced complexes with a constant activity ratio are concordant (Küken et al., 2022). Concordance partitions non-balanced complexes into concordance modules, which are in turn used to identify kinetic modules – subnetworks whose reactions are kinetically coupled(Langary et al., 2025).

The computational bottleneck in the study of the modularity of metabolic networks based on these concepts is concordance testing, which requires solving four linear programs (LPs) per pair of complexes, with number of pairs growing quadratically (e.g., reaching ≈10□for networks of ≈20,000 complexes after elementary-step expansion (Langary et al., 2025)). The reference MATLAB implementation (Küken et al., 2021, 2022; Langary et al., 2023, 2025) uses sampling-based candidate filtering but relies on a tensor representation that limits sample sizes (n□<□10) and solves pairs sequentially without transitivity pruning.

To address these limitations, we developed COCOA.jl, a Julia implementation that (1) solves LPs in parallel with cached JuMP models, (2) exploits the transitivity of concordance to skip already-implied pair checks, and (3) evaluates activity correlations in memory-efficient streaming batches. COCOA.jl builds on COBREXA.jl (Kratochvíl et al., 2025), so models from COBRApy, the COBRA Toolbox, or COBRA.jl can be used directly.

## 2 Implementation

### 2.1 Workflow

COCOA.jl exposes a four-step pipeline:

#### (i) Model preparation

Blocked reactions and orphan metabolites are removed by *prepare_model_for_concordance*. Because the concentration-robustness arguments of Langary *et al*. (2025) assume mass-action kinetics, *split_into_elementary_reactions* optionally expands each enzymatic reaction into elementary steps with either an ordered or a random enzyme-binding mechanism (Khodayari and Maranas, 2016; Liebermeister and Klipp, 2006; Segel, 1993).

#### (ii) Balanced complexes and trivial dependencies

The core function *activity_concordance_analysis* builds the stoichiometric constraints (reversible reactions can be split via *concordance_constraints*), identifies trivially balanced complexes (Küken et al., 2021) as well as trivially concordant pairs (Küken et al., 2022), and determines the sign of each complex’s activity through activity variability analysis. Activity variability analysis maximises and minimises each complex’s activity over the flux polytope, classifying it as strictly positive, strictly negative, or sign-variable. The sign classification reduces the number of Charnes–Cooper-transformed LPs that have to be solved per non-trivial pair.

#### (iii) Sampling-driven candidate filtering

Activities of complexes are sampled using artificial-centering hit-and-run on the flux polytope (ref to the sampler). A streaming filter retains only candidate pairs whose empirical activity-ratio coefficient of variation is below a user-defined threshold (default CV ≤ 0.01). The retained candidates are tested for concordance by solving the fractional linear programs that compute the minimum and maximum activity ratios; confirmed concordant pairs are merged by union–find, exploiting the equivalence-relation property of concordance to partition complexes into maximal mutually concordant sets. Pairs whose ratio range is non-singleton are added to a non-concordance set so that further candidates involving them can be skipped.

#### (iv) Kinetic modules and robustness

Kinetic modules are derived from concordance modules using the four-phase upstream algorithm of Langary *et al*. (2025): starting from extended modules (the concordance module augmented with the balanced complexes), complexes that violate autonomy (Phase I) or the feeding property (Phases II–IV) are iteratively removed; overlapping upstream sets are merged into the final kinetic modules. Within each kinetic module COCOA.jl then detects (a) metabolites with absolute concentration robustness and (b) metabolite pairs with absolute concentration ratio robustness (Langary *et al*., 2025) using differences in complexes participating in a kinetic module.

### 2.2 Algorithmic optimizations

The three optimizations together reduce the effective complexity of concordance analysis from quadratic to near-linear in the number of complexes (Fig. 1). *Parallel LP solution with cached JuMP models* eliminates the per-pair model-construction overhead that dominates the naive implementation. *Dynamic transitivity filtering* propagates already-confirmed concordance/non-concordance relations to prune pair checks on the fly. *Streaming computation* evaluates activity correlations in bounded-size batches, keeping peak memory tractable on networks with >10^4^ complexes. Across the benchmark dataset (Section 3) the candidate filter eliminated >99% of complex pairs before any LP was solved.

**Fig. 1.**
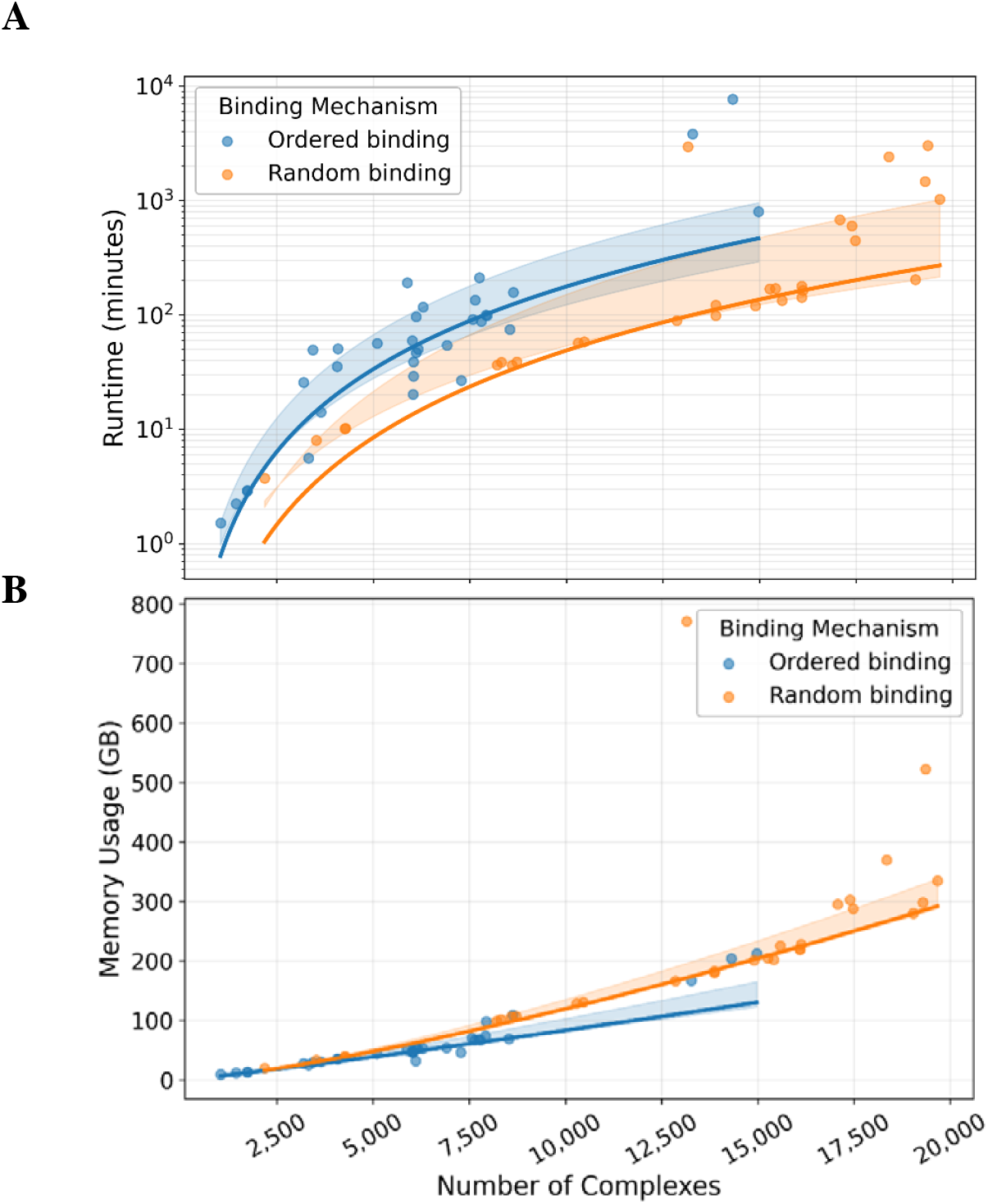
Scaling of required computational resources. (A) End-to-end COCOA.jl runtime for concordance and kinetic module analysis, log-scaled; (B) peak resident memory as a function of the number of complexes after elementary-step expansion, across the 62 benchmark runs (33 ordered-binding and 29 random-binding models). All runs used 64 CPU cores on an x86_64 Slurm cluster.

## 3 Benchmark against the reference implementation

### 3.1 Dataset and protocol

We benchmarked COCOA.jl against the MATLAB Upstream_Algorithm pipeline (github.com/ankueken/Upstream_Algorithm) on the 33 genome-scale models analysed by Langary *et al*. (2025). For each model the two enzyme-binding variants (ordered and random) were considered, giving 62 (model, variant) runs in total (three models yielded only an ordered-binding variant for the original reference implementation because of its random-variant runtime). All runs used the same pre-balanced elementary-step.mat files as the reference pipeline, so inputs were byte-identical. COCOA.jl and the MATLAB reference were executed with seed 42, sample size 10,000, CV threshold 0.01, and concordance tolerance 10^−2^; MATLAB output counts were independently cross-checked against Table S1 of Langary *et al*. (2025). Runs were performed on a shared Slurm cluster with 64 cores and up to 600 GB of memory per job.

### 3.2 Agreement with the reference implementation

For each model variant, we compared COCOA.jl and the reference implementation on three quantities highlighted by Langary *et al*. (2025): the size of the giant kinetic module (the largest kinetic module), the set of free metabolites with absolute concentration robustness (ACR), and the set of free metabolite pairs with absolute concentration-ratio robustness (ACRR). The per-model comparison is shown for ordered binding in Supplementary Fig. 1 and for random binding in Supplementary Fig. 2.

The giant kinetic module is of the same order of magnitude in COCOA.jl and in the reference implementation across the full dataset (Supplementary Fig. 1a and 2a). The sets of ACR free metabolites (Supplementary Fig. 1b, 2b) and ACRR free metabolite pairs (Supplementary Fig. 1c, 2c) track the reference on most networks; in case of differences, COCOA.jl tends to report additional. These differences likely arise from differences between the chosen LP solver (HiGHS in COCOA.jl vs. CPLEX in the reference), which can yield slightly different sets of concordant complex pairs and, consequently, different kinetic modules.

### 3.3 Runtime and memory scaling

We report runtime and peak memory for the concordance-analysis step only, because it is the performance-critical part of the pipeline: in every benchmark run concordance analysis accounted for more than 95% of the wall-clock time and for the peak memory footprint. COCOA.jl completed concordance analysis for every (model, variant) pair in the benchmark, including networks with up to 19,667 complexes after elementary-step expansion. Median concordance wall-clock time was 54 min for ordered binding and 134 min for random binding (Fig. 1a); median peak memory was 48 GB and 202 GB, respectively (Fig. 1b). The higher random-binding footprint reflects the denser connectivity of random-binding elementary-step networks. The reference MATLAB pipeline was unable to complete within the 600 GB / 48 h Slurm allocation on the three largest random-binding networks (*iAM_Pf480, iAM_Pk459, iAM_Pv461*, each >17,000 complexes), which is the main practical motivation for COCOA.jl.

## 4 Conclusion

COCOA.jl is a scalable, open-source Julia package that brings balanced-complex identification, concordance analysis, and kinetic module identification alongside downstream mining for identification of ACR and ACRR properties to genome-scale biochemical networks. It reproduces the published results of Langary *et al*. (2025) on the ordered-binding elementary-step decompositions of 33 genome-scale models at the level of reaction counts, complex counts, and kinetic module counts, and extends the analysis to random-binding variants and to networks that were previously intractable. Its integration with the COBREXA.jl ecosystem and its interoperability with Python and MATLAB should make it immediately usable in existing constraint-based workflows.

## Acknowledgements

We thank Dr. Damoun Langary for helpful discussions.

## Funding

The work was supported by funding from the University of Potsdam’s Research Focus Group “Evolutionary Systems Biology” and by the Deutsche Forschungsgemeinschaft (DFG, German Research Foundation) – SFB 1644/1 – 512328399.

## Conflict of interest

None declared.

## Notes

https://github.com/antoniofranky/COCOA.jl

### Competing Interest Statement

The authors have declared no competing interest.

