## Supplementary Figures for "COCOA.jl – A Julia package for high-performance analysis of concordance and kinetic modules in biochemical networks"

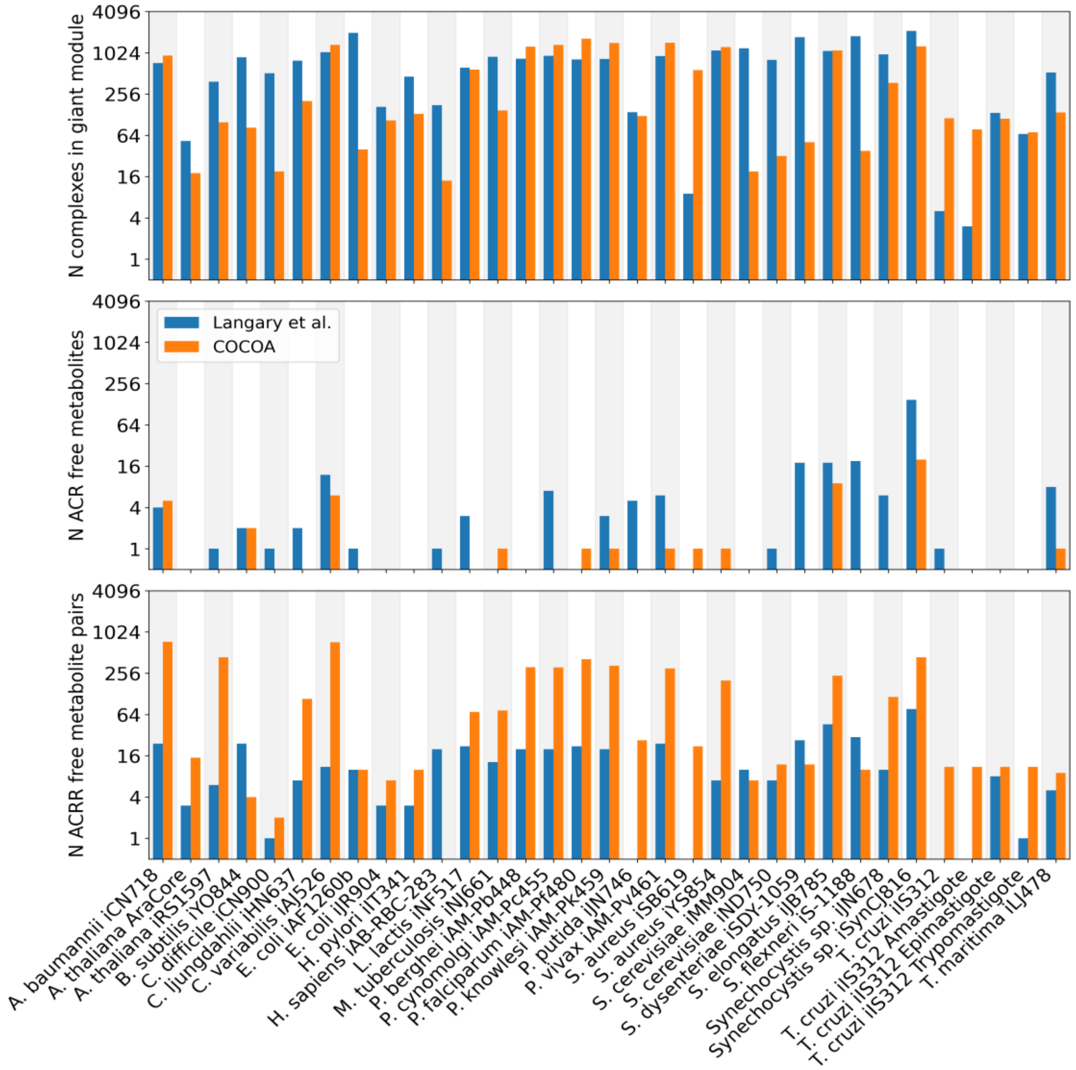


Supplementary Fig. 1. **Comparison of properties based on kinetic modules for ordered binding models.** Shown are the giant kinetic module size (top), the number of ACR free metabolites (middle) and the number of ACRR free metabolite pairs (bottom) for the 33 genome-scale models of Langary et al. (2025) under ordered-binding elementary-step expansion. Blue: reference MATLAB implementation (counts match Table S1 of the cited paper exactly); orange: COCOA.jl. Y-axis is log4-scaled to accommodate the three orders of magnitude spanned by the giant-module sizes.

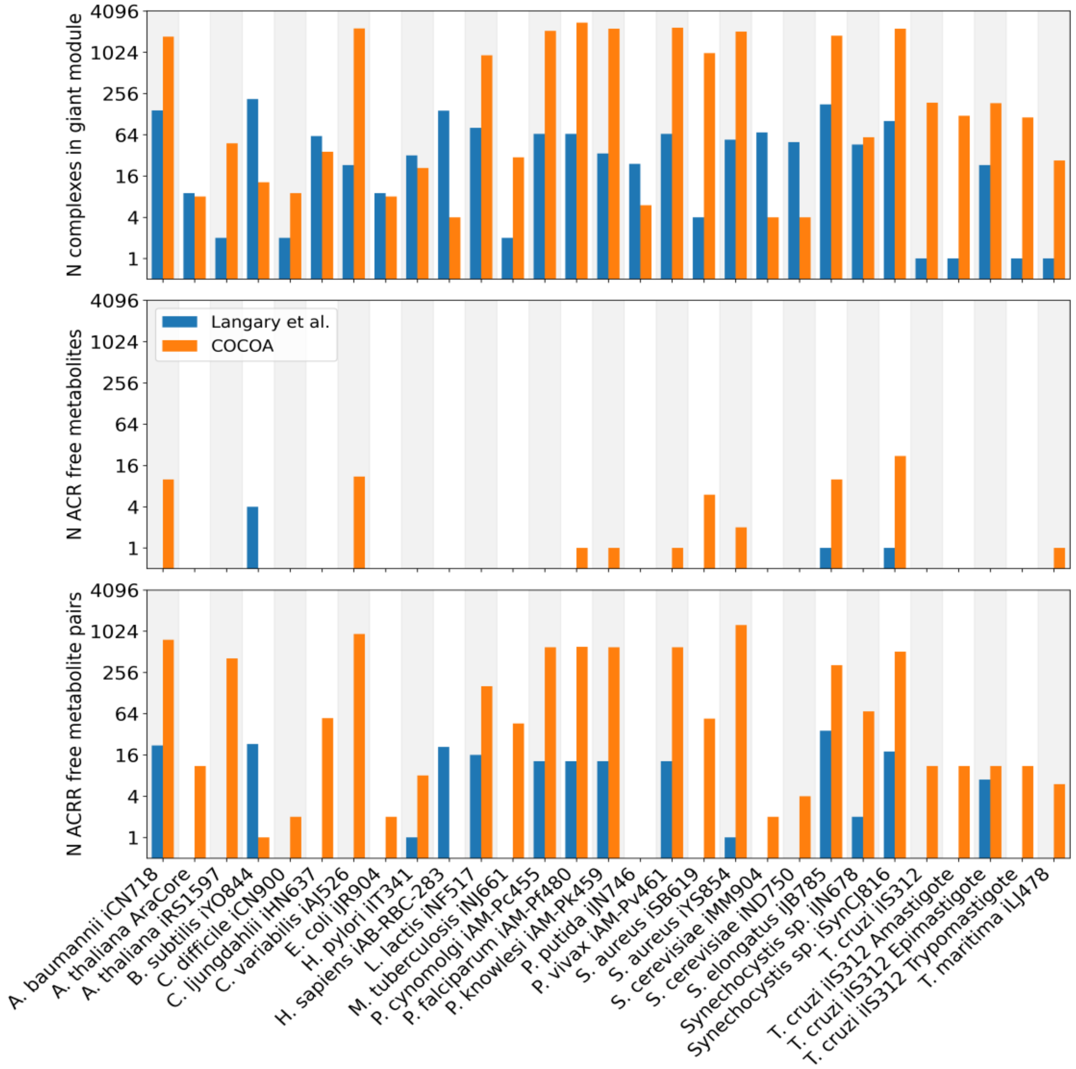


Supplementary Fig. 2. **Comparison of properties based on kinetic modules for random binding models.** The giant kinetic module size (top), the number of ACR free metabolites (middle) and the number of ACRR free metabolite pairs (bottom) for 29 genome-scale models of Langary et al. (2025) under random-binding elementary-step decomposition. Blue: reference MATLAB implementation; orange: COCOA.jl. Y-axis is log4-scaled to accommodate the three orders of magnitude spanned by the giant-module sizes.
